# Improved Modeling of Peptide-Protein Binding through Global Docking and Accelerated Molecular Dynamics Simulations

**DOI:** 10.1101/743773

**Authors:** Jinan Wang, Andrey Alekseenko, Dima Kozakov, Yinglong Miao

## Abstract

Peptides mediate up to 40% of known protein-protein interactions in higher eukaryotes and play a key role in cellular signaling, protein trafficking, immunology and oncology. However, it is challenging to predict peptide-protein binding with conventional computational modeling approaches, due to slow dynamics and high peptide flexibility. Here, we present a prototype of the approach which combines global peptide docking using *ClusPro PeptiDock* and all-atom enhanced simulations using Gaussian accelerated molecular dynamics (GaMD). For three distinct model peptides, the lowest backbone root-mean-square deviations (RMSDs) of their bound conformations relative to X-ray structures obtained from *PeptiDock* were 3.3 Å – 4.8 Å, being medium quality predictions according to the Critical Assessment of PRediction of Interactions (CAPRI) criteria. *GaMD* simulations refined the peptide-protein complex structures with significantly reduced peptide backbone RMSDs of 0.6 Å – 2.7 Å, yielding two high quality (sub-angstrom) and one medium quality models. Furthermore, the *GaMD* simulations identified important low-energy conformational states and revealed the mechanism of peptide binding to the target proteins. Therefore, *PeptiDock+GaMD* is a promising approach for exploring peptide-protein interactions.

## Introduction

Peptides mediate up to 40% of known protein-protein interactions in higher eukaryotes. Peptide binding plays a key role in cellular signaling, protein trafficking, immune response and oncology(Petsalaki and Russell, 2008;Das et al., 2013). In addition, peptides have served as promising drug candidates with high specificity and relatively low toxicity (Ahrens et al., 2012;Fosgerau and Hoffmann, 2015;Kahler et al., 2018;Lee et al., 2019). The number of peptide-based drugs being marketed is increasing in recent years (Ahrens et al., 2012;Fosgerau and Hoffmann, 2015;Kahler et al., 2018;Lee et al., 2019). Therefore, understanding the molecular mechanism of peptide-protein interactions is important in both basic biology and applied medical research.

Rational design of peptide-derived drugs usually requires structural characterization of the peptide-protein complexes. X-ray crystallography and nuclear magnetic resonance (NMR) have been utilized to determine high-resolution structures of peptide-protein complexes. These structures are often deposited into the Protein Data Bank (PDB) and also collected in specific databases focused on peptide-protein complex structures, including the PeptiDB(London et al., 2010), PepX(Vanhee et al., 2010) and PepBind(Das et al., 2013). Particularly, PeptiDB is a set of 103 non-redundant protein-peptide structures extracted from the PDB. The peptides are mostly 5-15 residues long (London et al., 2010). PepX contains 1431 non-redundant X-ray structures clustered based on the binding interfaces and backbone variations. There are 505 unique peptide-protein interfaces, including those for the major histocompatibility complex (MHC) (14%), thrombins (12%), α-ligand binding domains (8%), protein kinase A (5%), proteases and SH3 domains(Vanhee et al., 2010). The PepBind contains a comprehensive dataset of 3100 available peptide-protein structures from the PDB, irrespective of the structure determination methods and similarity in their protein backbone. More than 40% of the structures in PepBind are involved in cell regulatory pathways, nearly 20% in the immune system and ∼30% with protease or other hydrolase activities (Das et al., 2013). These databases have greatly facilitated structure-based modeling and drug design of peptide-protein interactions. However, the number of currently resolved structures is only a small fraction of the peptide-protein complexes, as limited by the difficulties and high cost of X-ray and NMR experiments.

Computational methods have been developed for predicting the peptide-protein complex structures. In this regard, modeling of peptide binding to proteins has been shown to be distinct from that of extensively studied protein-ligand binding and protein-protein interactions. Notably, small-molecule ligands are able to bind deeply buried sites in proteins, but peptides normally bind to the protein surface, especially in the largest pockets. On the other hand, protein partners usually have well defined 3D structures before forming protein-protein complexes, despite possible conformational changes during association. In contrast, most peptides do not have stable structures before forming complexes with proteins (Petsalaki and Russell, 2008). The biggest and immediate challenge for modeling of peptide-protein binding is that peptide structures are not known *a priori*. Furthermore, peptide-mediated interactions are often transient. The affinity of peptide-protein interactions is typically weaker than that of protein-protein interactions, because of the smaller interface between peptides and their protein partners. Therefore, new and robust computational approaches are developed to address the above challenges in the modeling of peptide-protein binding.

Molecular docking has proven useful in predictions of peptide-protein complex conformations (Ciemny et al., 2018). The commonly used approaches include template-based docking such as GalaxyPepDock(Lee et al., 2015), local docking of peptides to pre-defined binding sites such as Rosetta FlexPepDock(Raveh et al., 2011), HADDOCK(Trellet et al., 2013) and MDockPep(Xu et al., 2018), and global docking of free peptide binding to proteins such as CABS-dock (Kurcinski et al., 2015), PIPER-FlexPepDock (Alam et al., 2017) and PeptiDock (Alam et al., 2017). The template-based docking is highly efficient, but often limited to the availability of templates (Lee et al., 2015). Local docking is able to generate good quality models that meet the Critical Assessment of PRediction of Interactions (CAPRI) criteria (Janin et al., 2003). However, it requires *a priori* knowledge of the peptide binding site on the protein surface. In comparison, global peptide docking provides sampling of peptide binding over the entire protein surface without the need for pre-defined binding sites, but it is challenging to account for the system flexibility. In this regard, ClusPro *PeptiDock* has been developed for docking of motifs (short sequences) of peptides, which are found to sample only a small ensemble of different conformations (Alam et al., 2017). Structural ensemble of a peptide motif is built by retrieving motif structures from PDB that are very similar to the peptide’s bound conformation. A Fast-Fourier Transform (FFT) based docking is then used to quickly perform global rigid body docking of these fragments to the protein. *PeptiDock* is thus able to alleviate the peptide flexibility problem through ensemble docking of the peptide motifs. Nevertheless, it remains challenging to account for the high flexibility of the peptides. Overall, peptide docking often generates poor predictions that require further refinement to obtain CAPRI-quality models.

Molecular dynamics (MD) is a powerful technique that enables all-atom simulations of biomolecules. MD simulations are able to fully account for the flexibility of peptides and proteins during their binding (Knapp et al., 2015;Wan et al., 2015;Salmaso et al., 2017;Yadahalli et al., 2017;Kahler et al., 2018). MD has been used to refine binding poses of peptides in proteins in the pepATTRACT(de Vries et al., 2017) and AnchorDock(Ben-Shimon and Niv, 2015) docking protocols. However, it is challenging to sufficiently sample peptide-protein interactions through conventional MD (cMD) simulations, due to the slow dynamics and limited simulation timescales. Computational approaches that combine many cMD simulations provide improved sampling of peptide-protein interactions, including supervised MD (Salmaso et al., 2017) and weighted ensemble (Zwier et al., 2016). Notably, weighted ensemble of a total amount of ∼120 μs MD simulations has been obtained to investigate binding of an intrinsically disordered p53 peptide to the MDM2 Protein (Zwier et al., 2016). The simulation predicted binding rate constant agrees very well with the experiments. However, expensive computational resources would be needed for applications of cMD simulations in large-scale predictions of peptide-protein complex structures.

On the other hand, enhanced sampling MD methods have been developed to improve biomolecular simulations (Christen and van Gunsteren, 2008;Gao et al., 2008;Liwo et al., 2008;Dellago and Bolhuis, 2009;Abrams and Bussi, 2014;Spiwok et al., 2015;Miao and McCammon, 2016). Multi-ensemble Markov models (Paul et al., 2017), which combine cMD with Hamiltonian replica exchange enhanced sampling simulations, have been used to characterize peptide-protein binding and calculate kinetic rates of a nano-molar peptide inhibitor PMI to the MDM2 oncoprotein fragment (Paul et al., 2017). While cMD is able to simulate fast events such as peptide binding, enhanced sampling simulations can capture rare events such as peptide unbinding. The steered MD (Cuendet et al., 2011), temperature-accelerated MD (Lamothe and Malliavin, 2018) and MELD (Modeling by Employing Limited Data) using temperature and Hamiltonian replica exchange MD(Morrone et al., 2017) have also been applied to study peptide-protein binding. In comparison, more enhanced sampling methods have been applied in studies of protein-ligand binding and protein-protein interactions, including the umbrella sampling (Torrie and Valleau, 1977;Kastner, 2011;Rose et al., 2014), metadynamics (Laio and Parrinello, 2002;Alessandro and Francesco, 2008;Saleh et al., 2017a;Saleh et al., 2017b;Saleh et al., 2017c), adaptive biasing force (Darve and Pohorille, 2001;Darve et al., 2008), steered MD (Cuendet and Michielin, 2008;Gonzalez et al., 2011), replica exchange MD(Sugita and Okamoto, 1999;Okamoto, 2004), accelerated MD (aMD) (Hamelberg et al., 2004;Miao et al., 2015) and Gaussian accelerated MD (GaMD) (Miao et al., 2015;Miao and McCammon, 2017;Pang et al., 2017;Miao and McCammon, 2018). Overall, enhanced sampling simulations of peptide binding to proteins have been under explored. Peptide-protein binding shows distinct characteristics as described above and requires the development of improved enhanced sampling approaches.

Here, we present a prototype of a novel computational approach that combines global peptide docking using *PeptiDock* and all-atom enhanced sampling simulations using GaMD to model peptide-protein binding. Three model peptides have been selected from the PeptiDB database of non-redundant peptide-protein complex structures(London et al., 2010). They include peptide motifs “PAMPAR” (Peptide 1), “TIYAQV” (Peptide 2) and “RRRHPS” (Peptide 3), which bind to the SH3 domain, X-linked lymphoproliferative syndrome (XLP) protein SAP and human PIM1 kinase, respectively. Starting with the lowest RMSD conformation selected from top 10 models of *PeptiDock,* GaMD significantly refines the peptide-protein complex structures. Furthermore, the simulations provided important insights into the mechanism of peptide binding to target proteins at an atomistic level. Thus, *PeptiDock+GaMD* is a promising approach for exploring peptide-protein interactions.

## Methods

### A Computational Approach Combining *PeptiDock* and *GaMD*

A new computational approach was designed to predict peptide-protein complex structures by combining peptide docking with *PeptiDock* and all-atom enhanced sampling simulation with GaMD (**Figure S1** in Supplementary Material). Initial peptide-protein complex structures were obtained using the *ClusPro PeptiDock* server. The first step in the PeptiDock protocol is fragment search: the PDB database is searched for fragments containing the target peptide motif. The templates are clustered and an FFT-based rigid docking is applied to the cluster centroids. Top-scoring poses are clustered again and the centroids of the largest clusters are chosen as the final results (Porter et al., 2017). For the purpose of this study – to show the viability of the protocol – only one pose within top 10 models of *PeptiDock*, known to be near native, was selected for further refinement using GaMD simulations.

### System Setup

Three model peptides were selected from the PeptiDB database of non-redundant peptide-protein complex structures(London et al., 2010). They included peptide motifs “PAMPAR” (Peptide 1), “TIYAQV” (Peptide 2) and “RRRHPS” (Peptide 3), which bind to the SH3 domain, XLP protein and human PIM1 kinase, respectively. The free X-ray structures of target proteins is 1OOT, 1D1Z and 2J2I, respectively. The corresponding bound structures are 1SSH, 1D4T (Poy et al., 1999) and 2C3I (Pogacic et al., 2007), respectively. The free X-ray structures of the target proteins were used in the peptide docking and GaMD simulation. Both capped/neutral and uncapped/zwitterion terminus models were investigated in the GaMD simulations. In the neutral terminus model, the N- and C-termini were capped with ACE and NHE, respectively.

### Peptide Docking

The standard *ClusPro* PeptiDock protocol was used for all three systems. In the first step, receptor structures were specified: 1OOT chain A (Peptide 1), 1D1Z chain A (Peptide 2) and residues 125-305 of 2J2I chain B (Peptide 3). The next step was specifying motifs – the templates for searching fragments in PDB database. The motif was specified as subsequence of the peptide with one or more wildcard symbols. Wildcards could be of two forms: “X”, denoting any amino acid substitution, and “[…]”, denoting substitution by any amino acid from the list. *E.g.*, “[FT]” means that either Phe or Tyr can take this place. It is recommended to adjust the motif to yield between 100 and 1,000 hits, while preserving the essential features for binding. For the studied systems, the following motifs were used for fragment search: “PXMPXR” for Peptide 1 (107 hits, see Ref. (Hou et al., 2012)), “TI[YF]XX[VI]” for Peptide 2 (686 hits, see Ref.(Poy et al., 1999)) and “RXRHXS” for Peptide 3 (198 hits, see Ref. (Bullock et al., 2005)). Since PDB contains bound structures of the studied systems, a number of PDB entries were explicitly excluded from template search, as listed in Table S4. The next steps were performed automatically by the server (Porter et al., 2017), being the same for all systems. The extracted fragments were changed to the target peptide sequence using backbone-dependent rotamer library(Dunbrack and Karplus, 1993). The extracted fragments (hits) were clustered using the greedy algorithm according to their pairwise root-mean-square deviation (RMSD), with 0.5 Å cluster radius. The centroids of top 25 clusters were docked to the receptor using rigid-body FFT docking(Kozakov et al., 2006), exhaustively sampling all possible mutual orientations of the receptor and ligand, and ranking them using a special scoring function with a mixture of physics-based and knowledge-based terms (Kozakov et al., 2006;Chuang et al., 2008).The top-scoring poses of each fragment were pooled together and clustered based on their pairwise RMSDs, with 3.5 Å cluster radius. The clusters were ranked according to their sizes (Kozakov et al., 2005). The centroids of ten largest clusters were subjected to energy minimization with a CHARMM19-based force field using the ABNR algorithm. To demonstrate the protocol, only the lowest RMSD conformation obtained from top 10 PeptiDock models of each peptide was selected for refinement using GaMD simulations. The ranks of docking poses with the lowest peptide backbone RMSDs used were 9, 5 and 10 for Peptides 1, 2 and 3, respectively. It is important to note that each of the top-10 docking models will be refined and scored in a full version of the protocol in further studies.

### GaMD Enhanced Sampling Simulations

GaMD was applied to refine the peptide-protein complex structures. Complexes were solvated in explicit water using *tleap* from the AMBER 18 package (Case et al., 2018). The Na^+^ and Cl^−^ ions were added to neutralize the system charge. The AMBER ff14SB force field parameters (Maier et al., 2015) and TIP3P model (Jorgensen et al., 1983) were used for the proteins/peptides and water molecules, respectively. Each system was minimized using steepest descent for 50000 steps and conjugate gradient for another 50000 steps. After minimization, the system was heated from 0 to 310 K in 1 ns simulation by applying 1 kcal/(mol•Å^2^) harmonic position restraints to the protein and peptide heavy atoms with a constant number, volume and temperature (NVT) ensemble. Each system was further equilibrated using a constant number, pressure and temperature (NPT) ensemble at 1 atm and 310 K for 1ns with same restraints as in the NVT run. Another 2ns cMD simulations were performed to collect potential energy statistics (including the maximum, minimum, average and standard deviation). Then 18 ns GaMD equilibration after applying the boost potential was performed. Finally, four independent 300 *ns* GaMD production runs with randomized initial atomic velocities were performed on each peptide system. Simulation frames were saved every 0.2 ps for analysis. Snapshots of all four GaMD productions (1,200 ns in total) were combined for clustering to identify peptide binding conformations using the hierarchical agglomerative algorithm in CPPTRAJ (Roe and Cheatham, 2013). The cutoff was set to 3.5 Å for the peptide backbone RMSD to form a cluster. The *PyReweighting* toolkit (Miao et al., 2014) was applied to reweight GaMD simulations and recover the original free energy or potential of mean force (PMF) profiles of the peptide-protein systems. The RMSDs of the peptide and protein backbone were used as reaction coordinates. Detailed descriptions of GaMD theory and energetic reweighting were shown in supplementary material.

## Results

### Prediction of Peptide Binding Conformations through Docking and GaMD Simulations

There were no significant conformational changes in the protein during binding of Peptides 1 and 3 (**Figure 1A and 1C**). In comparison, binding of Peptide 2 induced a large structural rearrangement of the loop involving residues 67 to 74 in the protein (**Figure 1B**). In addition, Peptide 3 is highly charged as its first three N-terminal residues in the sequence are all arginine. These features of Peptides 2 and 3 raised the difficulty in accurate prediction of their peptide-protein complex structures. Peptide docking with *PeptiDock* showed different levels of accuracy: RMSDs of the peptide backbone compared with the bound X-ray structures were 3.3 Å, 3.5 Å and 4.8 Å for the three peptides, respectively (**Figures 1A-1C** and **Table 1**). They were all medium quality predictions according to the CAPRI criteria (Janin et al., 2003).

**Table 1.**
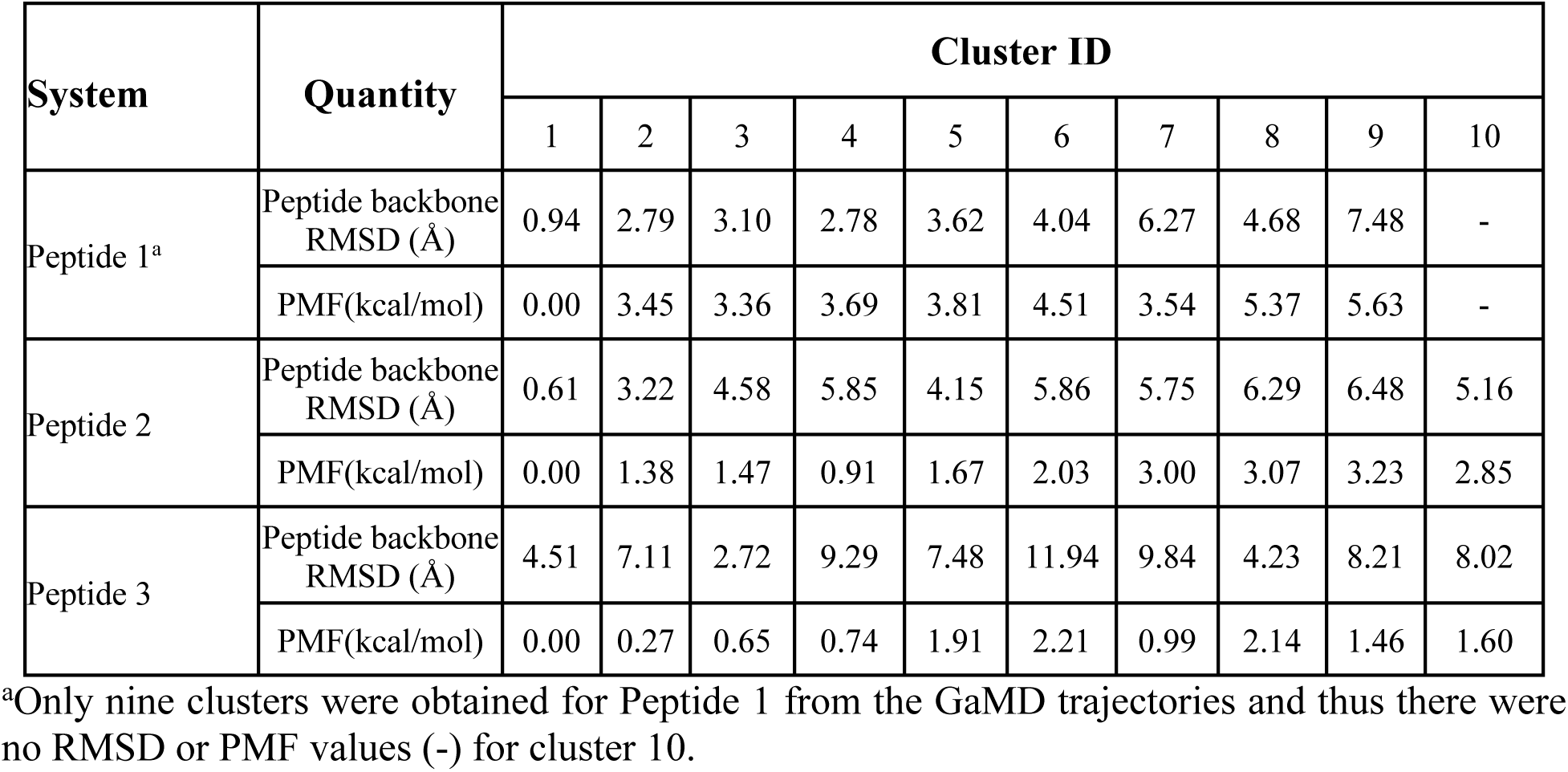
Comparison of 10 top-ranked clusters of three model peptides using the *PeptiDock+GaMD* approac

| System | Quantity | Cluster ID |  |  |  |  |  |  |  |  |  |
| --- | --- | --- | --- | --- | --- | --- | --- | --- | --- | --- | --- |
|  |  | 1 | 2 | 3 | 4 | 5 | 6 | 7 | 8 | 9 | 10 |
| Peptide 1<sup>a</sup> | Peptide backbone RMSD (Å) | 0.94 | 2.79 | 3.10 | 2.78 | 3.62 | 4.04 | 6.27 | 4.68 | 7.48 | - |
|  | PMF(kcal/mol) | 0.00 | 3.45 | 3.36 | 3.69 | 3.81 | 4.51 | 3.54 | 5.37 | 5.63 | - |
| Peptide 2 | Peptide backbone RMSD (Å) | 0.61 | 3.22 | 4.58 | 5.85 | 4.15 | 5.86 | 5.75 | 6.29 | 6.48 | 5.16 |
|  | PMF(kcal/mol) | 0.00 | 1.38 | 1.47 | 0.91 | 1.67 | 2.03 | 3.00 | 3.07 | 3.23 | 2.85 |
| Peptide 3 | Peptide backbone RMSD (Å) | 4.51 | 7.11 | 2.72 | 9.29 | 7.48 | 11.94 | 9.84 | 4.23 | 8.21 | 8.02 |
|  | PMF(kcal/mol) | 0.00 | 0.27 | 0.65 | 0.74 | 1.91 | 2.21 | 0.99 | 2.14 | 1.46 | 1.60 |

<sup>a</sup>Only nine clusters were obtained for Peptide 1 from the GaMD trajectories and thus there were no RMSD or PMF values (-) for cluster 10.

**Figure 1.**
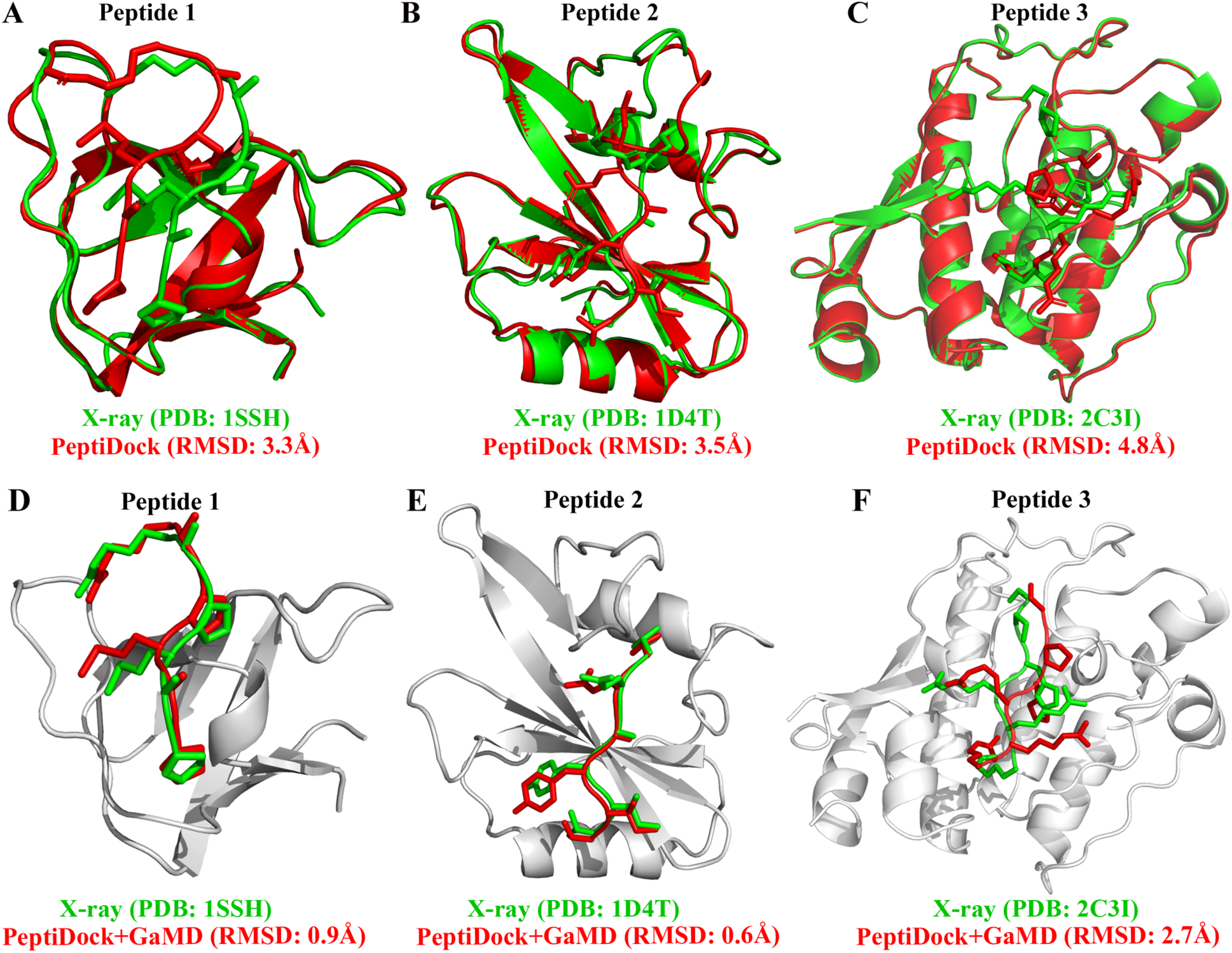
Docking poses (red) of three peptide motifs obtained using *PeptiDock* are compared with X-ray structures (green): (A) Peptide 1 “PAMPAR”, (B) Peptide 2 “TIYAQV” and (C) Peptide 3 “RRRHPS”; Binding poses (red) of three model peptides obtained using the “*PeptiDock+GaMD”* are compared with X-ray structures (green): (D) Peptide 1, (E) Peptide 2 and (F) Peptide 3.

Next, GaMD simulations were performed to refine the docking models. Analysis of simulation trajectories showed that the GaMD simulations were able to effectively refine the peptide binding pose. For Peptides 1 and 2, RMSDs of the peptide backbone relative to the X-ray structures decreased to < 1 Å during the GaMD simulations (**Figure 2A and 2B**). Peptide 1 bound tightly to the protein target site throughout the four GaMD simulations. Peptide 2 reached the native conformation within ∼10 ns, ∼90 ns, ∼120 ns and ∼170 ns in the four GaMD simulations and stayed tightly bound during the remainder of the simulations. In comparison, Peptide 3 exhibited higher fluctuations and sampled the near-native conformation transiently during the GaMD simulations (**Figure 2C**). Nevertheless, the minimum RMSDs of peptide backbone compared with X-ray structures were identified to be 0.20 Å, 0.22 Å and 0.73 Å for the three peptides, respectively (**Figures 2A-2C**).

**Figure 2.**
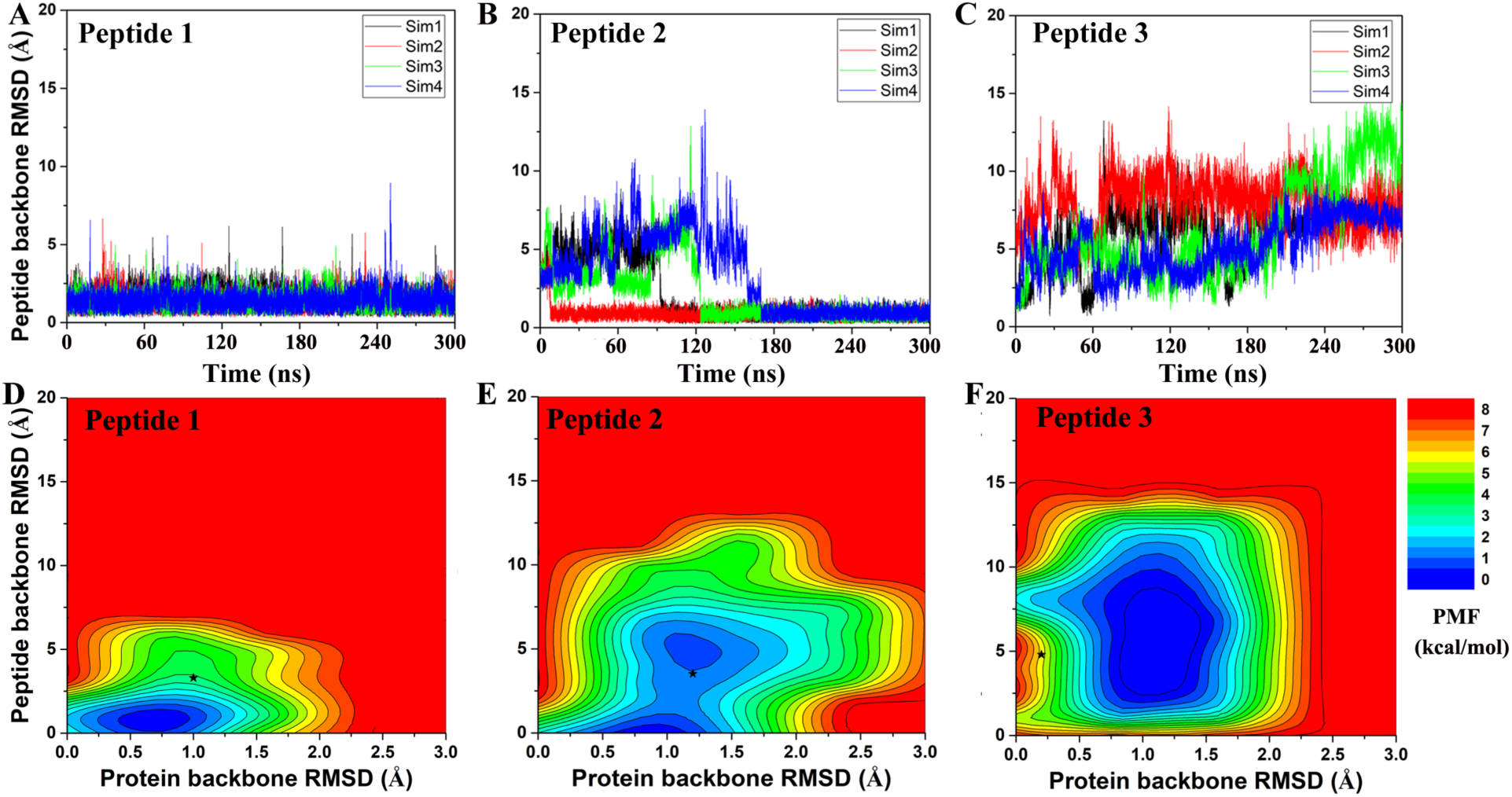
Time courses of peptide backbone RMSD obtained from four 300*ns* GaMD simulations on (A) Peptide 1, (B) Peptide 2 and (C) Peptide 3. 2D potential of mean force (PMF) regarding the peptide backbone RMSD and protein backbone RMSD for (D) Peptide 1, (E) Peptide 2 and (F) Peptide 3. The black stars indicate the initial binding poses obtained using *PeptiDock*.

Furthermore, GaMD simulation snapshots of the peptide conformations were clustered using the backbone RMSDs relative to the X-ray structures. This procedure was similar to analysis of the peptide docking poses. The 10 top-ranked clusters of peptide conformations with the lowest free energies were obtained. The 1^st^ top-ranked cluster exhibited peptide backbone RMSDs of 0.94 Å and 0.61 Å for Peptides 1 and 2, respectively (**Figures 1D-1E** and **Table 1**). For Peptide 3, the 3^rd^ top-ranked cluster showed the smallest peptide backbone RMSD of 2.72 Å (**Figure 1F** and **Table 1**). According to the CAPRI criteria (Janin et al., 2003), structural predictions for Peptides 1 and 2 were of sub-angstrom high quality and medium quality for Peptide 3. Therefore, GaMD simulations significantly refined docking conformations of the three peptide-protein complex structures. The simulation predicted bound conformations of the peptides were in excellent agreement with experimental X-ray structures with 0.6 Å – 2.7 Å in the peptide backbone RMSDs. In comparison, docking poses of the three peptides obtained from *PeptiDock* showed RMSDs of 3.3 Å – 4.8 Å (**Table 1**).

### Peptide Binding Mechanism Revealed from *GaMD*

Free energy profiles were calculated from the GaMD simulations using the protein and peptide backbone RMSDs relative to the bound X-ray structures as reaction coordinates. For Peptide 1, only one low-energy minimum was identified near the native bound state (**Figure 2D**). This was consistent with the clustering result that the peptide backbone RMSD of the 1^st^ top-ranked cluster was only 0.9Å.

For peptide 2, two low-energy minima were identified, corresponding to peptide backbone RMSDs of 0.5 Å and 4.2 Å, respectively (**Figure 2E**). As described above, the binding of Peptide 2 induced a significant conformational change in the protein loop of residues 67 to 74 (**Figure 1B**). Thus, the loop backbone RMSD and peptide backbone RMSD relative to the bound X-ray structure were also used as reaction coordinates to compute another two-dimensional free energy profile (**Figure 3A)**. The protein loop was highly flexible, sampling a large conformational space. The loop backbone RMSD ranged from ∼0.2 Å to ∼8.0 Å. This loop sampled two low-energy conformations, including the “Open” (bound) (RMSD < 1 Å) and “Closed” (free) states (RMSD ∼3-6 Å) (**Figure 3**). Compared to the “Open” state, the “Closed” loop moved closer to the core domain of protein (**Figure 3B**). GaMD simulations successfully captured the conformational change of this loop. The peptide and protein loop accommodated each other to form the final bound conformation (**Figure 3**), suggesting an “induced fit” mechanism.

**Figure 3.**
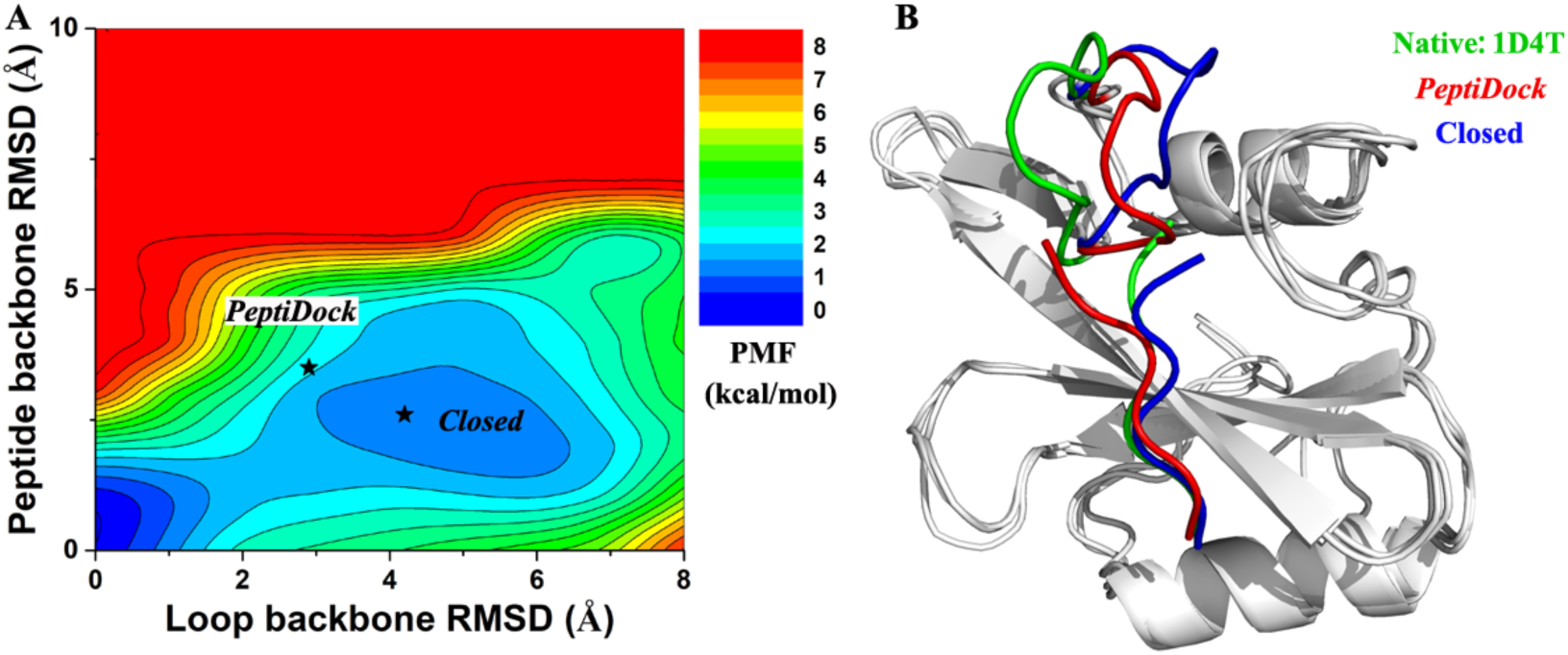
(A) 2D PMF calculated for binding of Peptide 2 regarding RMSDs of the peptide backbone and protein loop (residues 67 to 74) relative to the X-ray structure (PDB: 1D4T). (B) Representative conformation of “Closed” state (blue) in compared with initial conformation from “*PeptiDock*” (red) and X-ray structure (green).

For Peptide 3, GaMD sampled a broad low-energy well, centered at the ∼4.3 Å and ∼1.0 Å RMSDs for the peptide and protein backbone relative to the bound X-ray structure (**Figure 2F**). Overall, this peptide-protein complex underwent high fluctuations, visiting a large conformational space. Nevertheless, GaMD simulations sampled the native binding pose of Peptide 3, for which the peptide backbone RMSD decreased to ∼1 Å at ∼60ns and 160 ns during one of the GaMD production runs (Sim1) (**Figure 2C**). In contrast to binding of Peptide 2 that involved induced fit of the protein receptor, binding of Peptides 1 and 3 did not induce significant conformational change of the receptors.

### Effects of the Terminal Residue Charges on Peptide Binding

In addition to the neutral terminus model as described above, we simulated another model of the three peptides with zwitterion terminal residues that were charged. Compared with the neutral terminus models, larger fluctuations were observed in the zwitterion terminus models of the three peptides (**Figures S2-S4**). For Peptides 2 and 3, their backbone RMSDs could reach large values of ∼40 Å and ∼20 Å, respectively. These results suggested that the peptides could dissociate from the initial near-native bound pose obtained from docking. Furthermore, 10 top-ranked clusters of peptide conformations with the lowest free energies were also calculated through structural clustering and energetic reweighting (see Methods for details). For Peptide 1, the 1^st^ top-ranked cluster exhibited the smallest backbone RMSD of 1.22 Å relative to the X-ray structure (**Figure S5A** and **Table S1**). The 2^nd^ top-ranked clusters exhibited the smallest backbone RMSDs of 0.62 Å and 3.88 Å for Peptides 2 and 3, respectively (**Figures S5B-S5C and Tables S2-S3**). In summary, peptides with zwitterion terminal residues underwent higher fluctuations and the simulation predicted bound conformations deviated more from the native X-ray structures compared with the neutral terminal models.

## Discussion

We have demonstrated that *GaMD* can successfully refined *PeptiDock* docking poses, and thus established the possibility of *PeptiDok+GaMD* combination to predict peptide-protein complex structure and explore the peptide binding mechanism. Three peptides with different difficulty levels were selected as model systems. Peptide 1 was the easiest one as the peptide is rigid and there was no conformational change in the protein during peptide binding. Both Peptides 2 and 3 were challenging for predicting bound conformations accurately. The binding of Peptide 2 involved a significant structural rearrangement of the residue 67-74 loop in the protein. Peptide 3 with dense residue charges proved difficult for both docking and GaMD simulations. Nevertheless, the GaMD refinement achieved high quality models for both Peptides 1 and 2, and medium quality prediction for Peptide 3. This approach showed promise to be widely applicable for other peptide-protein binding systems.

It is difficult for the current docking programs to account for large conformational changes of proteins during peptide binding (Ciemny et al., 2018). Even in the flexible docking calculation, only movements of protein side chains are often taken into account. This raised a challenge in the modeling of Peptide 2. On the other hand, cMD simulations could account for flexibility of the peptide and protein and had been applied to refine docking poses of peptides in proteins (Ben-Shimon and Niv, 2015;de Vries et al., 2017). However, cMD could suffer from insufficient sampling and limited simulation timescales. Thus, the GaMD enhanced sampling method has been in this study. Remarkably, GaMD effectively captured the loop movement of Peptide 2 (**Figure 3**) and greatly refined the peptide docking poses (**Figure 1E**).

In summary, *PeptiDock+GaMD* has been demonstrated on predicting the peptide-protein complex structures and revealing important insights into the mechanism of peptide binding to proteins, using three distinct peptides as model systems. In the future, all top-10 models of the *ClusPro PeptiDock* will be refined with GaMD and a larger number of protein-peptide systems will be evaluated systematically. Furthermore, the effects of different force fields (e.g. CHARMM36) and solvent models (e.g., TIP4P and implicit solvent, etc.) (Kuzmanic et al., 2019) are to be further investigated. Development of novel protocols to increase the accuracy of peptide-protein structural prediction will facilitate peptide drug design. Advances in the computational methods and computing power are expected to help us to address these challenges.

## Author Contributions

Y.M. and D.K designed research; J.W. and A.A. performed research; J.W., A.A., D. K., and Y.M. analyzed data; and J.W., A.A., D. K., and Y.M. wrote the paper.

## Acknowledgements

This work was supported in part by the National Institutes of Health (R01GM132572), National Science Foundation (AF 1816314, DBI 1759277), Russian Science Foundation (19-74-00090), American Heart Association (Award 17SDG33370094) and the startup funding in the College of Liberal Arts and Sciences at the University of Kansas. Computing time was provided on the Comet and Stanford EXtream supercomputers through the Extreme Science and Engineering Discovery Environment award TG-MCB180049 and the Edison and Cori supercomputers through the National Energy Research Scientific Computing Center project M2874.

## Supplementary Material

The Supplementary Material for this article can be found online at https://www.frontiersin.org/.

