## Supplementary Material for "Improved Modeling of Peptide-Protein Binding through Global Docking and Accelerated Molecular Dynamics Simulations"

**Supplementary Methods**

**Gaussian accelerated Molecular Dynamics (GaMD)**

GaMD enhances the conformational sampling of biomolecules by adding a harmonic boost potential to reduce the system energy barriers (Miao et al., 2015). When the system potential $V(\vec{r})$ is lower than a reference energy *E*, the modified potential $V^*(\vec{r})$ of the system is calculated as:

$$V^*(\vec{r}) = V(\vec{r}) + \Delta V(\vec{r})$$

$$\Delta V(\vec{r}) = \begin{cases} \frac{1}{2}k\big(E - V(\vec{r})\big)^2, & V(\vec{r}) < E \\ 0, & V(\vec{r}) \geq E, \end{cases} \qquad \textbf{(1)}$$

Where k is the harmonic force constant. The two adjustable parameters *E* and *k* are automatically determined based on three enhanced sampling principles. First, for any two arbitrary potential values $V_1(\vec{r})$ and $V_2(\vec{r})$ found on the original energy surface, if $V_1(\vec{r}) < V_2(\vec{r})$, $\Delta V$ should be a monotonic function that does not change the relative order of the biased potential values; i.e., $V_1^*(\vec{r}) < V_2^*(\vec{r})$. Second, if $V_1(\vec{r}) < V_2(\vec{r})$, the potential difference observed on the smoothened energy surface should be smaller than that of the original; i.e., $V_2^*(\vec{r}) - V_1^*(\vec{r}) < V_2(\vec{r}) - V_1(\vec{r})$. By combining the first two criteria and plugging them in the formula of $V^*(\vec{r})$ and $\Delta V$, we obtain

$$V_{max} \leq E \leq V_{min} + \frac{1}{k}, \qquad \textbf{(2)}$$

Where $V_{min}$ and $V_{max}$ are the system minimum and maximum potential energies. To ensure that **Eq. 2** is valid, *k* has to satisfy: $k \leq 1/(V_{max} - V_{min})$. Let us define: $k = k_0 \cdot 1/(V_{max} - V_{min})$, then $0 < k_0 \leq 1$. Third, the standard deviation (SD) of $\Delta V$ needs to be small enough (i.e. a narrow distribution) to ensure accurate reweighting using cumulant expansion to the second order: $\sigma_{\Delta V} = k\big(E - V_{avg}\big)\sigma_V \leq \sigma_0$, where $V_{avg}$ and $\sigma_V$ are the average and SD of $\Delta V$ with $\sigma_0$ as a user-specified upper limit (e.g., $10k_BT$) for accurate reweighting. When $E$ is set to the lower bound $E = V_{max}$ according to **Eq. 2**, $k_0$ can be calculated as

$$k_0 = \min(1.0, k_0') = min\left(1.0, \frac{\sigma_0}{\sigma_V} \cdot \frac{V_{max}-V_{min}}{V_{max}-V_{avg}}\right), \quad \textbf{(3)}$$

Alternatively, when the threshold energy $E$ is set to its upper bound $E = V_{min} + 1/k$, $k_0$is set to:

$$k_0 = k_0'' \equiv \left(1 - \frac{\sigma_0}{\sigma_V}\right) \cdot \frac{V_{max}-V_{min}}{V_{avg}-V_{min}}, \quad \textbf{(4)}$$

If $k_0''$is calculated between 0 and 1. Otherwise, $k_0$is calculated using **Eq. 3**.

**Energetic Reweighting of GaMD Simulations**

To calculate potential of mean force (PMF), the probability distribution along a reaction coordinate is written as $p^*(A)$ . Given the boost potential $\Delta V(r)$ of each frame, $p^*(A)$ can be reweighted to recover the canonical ensemble distribution $p(A)$, as:

$$p(A_j) = p^*(A_j)\frac{\langle e^{\beta\Delta V(r)}\rangle_j}{\sum_{i=1}^{M}\langle p^*(A_i)e^{\beta\Delta V(r)}\rangle_i}, \ j = 1, \dots, M, \quad \textbf{(5)}$$

where $M$ is the number of bins, $\beta = k_BT$ and $\langle e^{\beta\Delta V(r)}\rangle_j$ is the ensemble-averaged Boltzmann factor of $\Delta V(r)$ for simulation frames found in the $j^{\text{th}}$ bin. The ensemble-averaged reweighting factor can be approximated using cumulant expansion:

$$\langle e^{\beta\Delta V(r)}\rangle = exp\left\{\sum_{k=1}^{\infty}\frac{\beta^k}{k!}C_k\right\}, \quad \textbf{(6)}$$

where the first two cumulants are given by:

$$\begin{gathered} C_1 = \langle\Delta V\rangle, \\ C_2 = \langle\Delta V^2\rangle - \langle\Delta V\rangle^2 = \sigma_v^2. \end{gathered} \quad \textbf{(7)}$$

The boost potential obtained from GaMD simulations usually follows near-Gaussian distribution (Miao and McCammon, 2017). Cumulant expansion to the second order thus provides a good approximation for computing the reweighting factor (Miao et al., 2014;Miao et al., 2015). The reweighted free energy $F(A) = -k_B T \ln p(A)$ is calculated as:

$$F(A) = F^*(A) - \sum_{k=1}^{2} \frac{\beta^k}{k!} C_k + F_c, \tag{8}$$

where $F^*(A) = -k_B T \ln p^*(A)$ is the modified free energy obtained from GaMD simulation and $F_c$ is a constant.

Table S1. Comparison of 10 top-ranked clusters of Peptide 1 with different terminus models using the *PeptiDock+GaMD* approach

| Cluster id | *PeptiDock+GaMD* *(Neutral terminus)* | | *PeptiDock+GaMD* *(Zwitterion terminus)* | |
|---|---|---|---|---|
| | Peptide backbone RMSD (Å) | PMF (kcal/mol) | Peptide backbone RMSD (Å) | PMF (kcal/mol) |
| 1 | 0.94 | 0.00 | 1.22 | 0.00 |
| 2 | 2.79 | 3.45 | 2.67 | 2.81 |
| 3 | 3.10 | 3.36 | 3.58 | 2.46 |
| 4 | 2.78 | 3.69 | 3.48 | 2.70 |
| 5 | 3.62 | 3.81 | 3.69 | 2.90 |
| 6 | 4.04 | 4.51 | 2.36 | 2.87 |
| 7 | 6.27 | 3.54 | 4.24 | 3.44 |
| 8 | 4.68 | 5.37 | 3.56 | 2.95 |
| 9 | 7.48 | 5.63 | 3.01 | 3.53 |
| 10[a] | - | - | 3.23 | 2.95 |

[a] Only nine clusters were obtained for Peptide 1 from the GaMD trajectories and thus there were no RMSD or PMF values (-) for cluster 10.

.

Table S2. Comparison of 10 top-ranked clusters of Peptide 2 with different terminus models using the *PeptiDock+GaMD* approach

| Cluster id | *PeptiDock+GaMD* (Neutral terminus) | | *PeptiDock+GaMD* (Zwitterion terminus) | |
|---|---|---|---|---|
| | Peptide backbone RMSD (Å) | PMF (kcal/mol) | Peptide backbone RMSD (Å) | PMF (kcal/mol) |
| 1 | 0.61 | 0.00 | 5.50 | 0.27 |
| 2 | 3.22 | 1.38 | 0.62 | 0.00 |
| 3 | 4.58 | 1.47 | 9.10 | 0.98 |
| 4 | 5.85 | 0.91 | 14.11 | 1.00 |
| 5 | 4.15 | 1.67 | 11.37 | 1.67 |
| 6 | 5.86 | 2.03 | 23.01 | 1.68 |
| 7 | 5.75 | 3.00 | 15.78 | 1.39 |
| 8 | 6.29 | 3.07 | 4.93 | 1.62 |
| 9 | 6.48 | 3.23 | 16.10 | 1.48 |
| 10 | 5.16 | 2.85 | 25.89 | 1.96 |

Table S3. Comparison of 10 top-ranked clusters of Peptide 3 with different terminus models using the *PeptiDock+GaMD* approach

| Cluster id | *PeptiDock+GaMD* (Neutral terminus) | | *PeptiDock+GaMD* (Zwitterion terminus) | |
|---|---|---|---|---|
| | Peptide backbone RMSD (Å) | PMF (kcal/mol) | Peptide backbone RMSD (Å) | PMF (kcal/mol) |
| 1 | 4.51 | 0.00 | 9.45 | 0.18 |
| 2 | 7.11 | 0.27 | 3.88 | 0.75 |
| 3 | 2.72 | 0.65 | 8.31 | 0.62 |
| 4 | 9.29 | 0.74 | 17.47 | 0.50 |
| 5 | 7.48 | 1.91 | 4.30 | 0.11 |
| 6 | 11.94 | 2.21 | 3.67 | 0.38 |
| 7 | 9.84 | 0.99 | 5.33 | 0.46 |
| 8 | 4.23 | 2.14 | 9.67 | 1.50 |
| 9 | 8.21 | 1.46 | 11.07 | 1.32 |
| 10 | 8.02 | 1.60 | 3.68 | 0.0 |

Table S4. Parameters of *ClusPro PeptiDock* runs

| Peptide # | Peptide sequence | Peptide motif | Excluded PDBs | Hits found |
|---|---|---|---|---|
| 1 | PAMPAR | PXMPXR | 1SSH 1Q2V 2B3H 2B3K 2B3L 2BER 2BPA 2BZD 2G6P 2GZ5 2J2F 2NQ6 2NQ7 2X6U 2X6V 2XPY 2XPZ 2XQ0 2XZ0 2XZ1 3T2B 3T2C 3T2D 3T2E 3T2F 3T2G | 107 |
| 2 | TIYAQV | TI[YF]XX[VI] | 1D4T 1D4W 1I3Z 1M27 | 686 |
| 3 | RRRHPS | RXRHXS | 2C3I 3CXW 3CY2 3CY3 4GW8 | 198 |

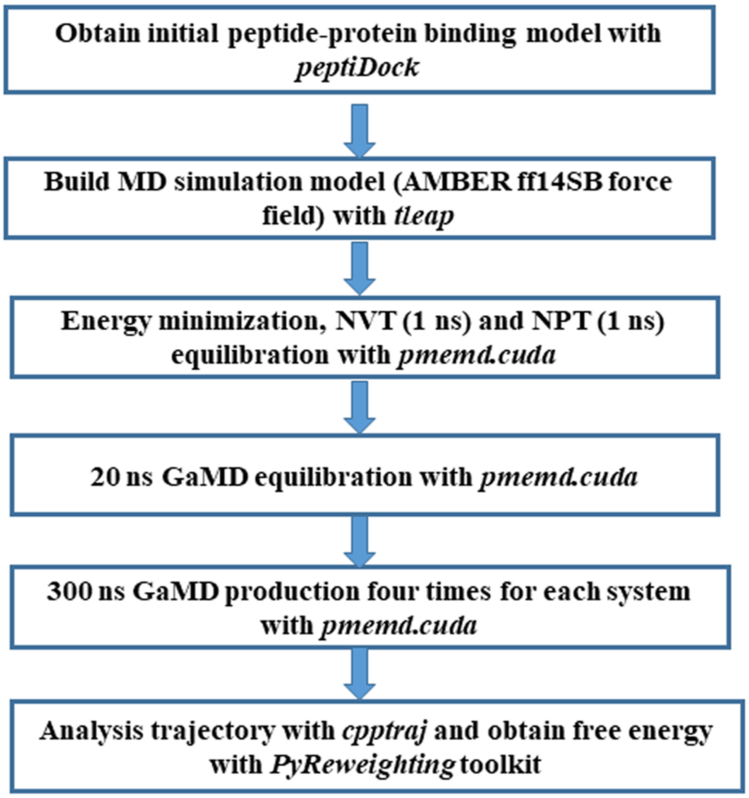


Figure S1. Workflow of the *PeptiDock+GaMD* approach

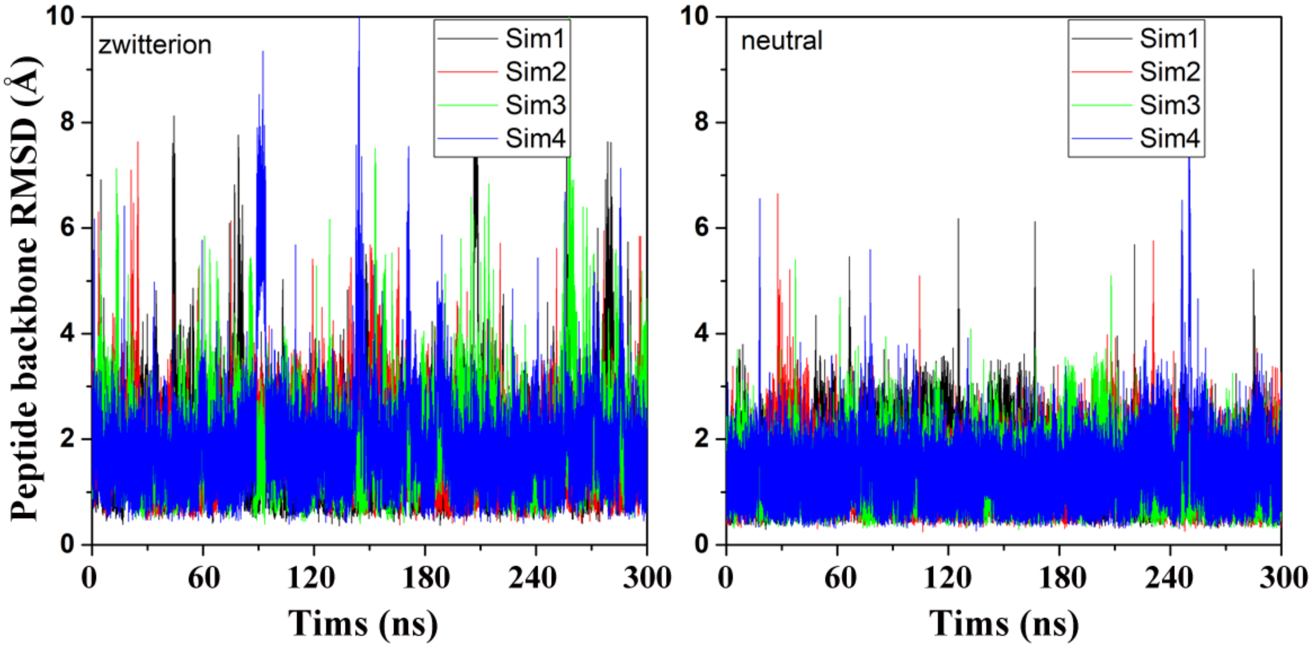


Figure S2. Comparison of zwitterion (left) and neutral (right) terminus models in GaMD simulations of Peptide 1.

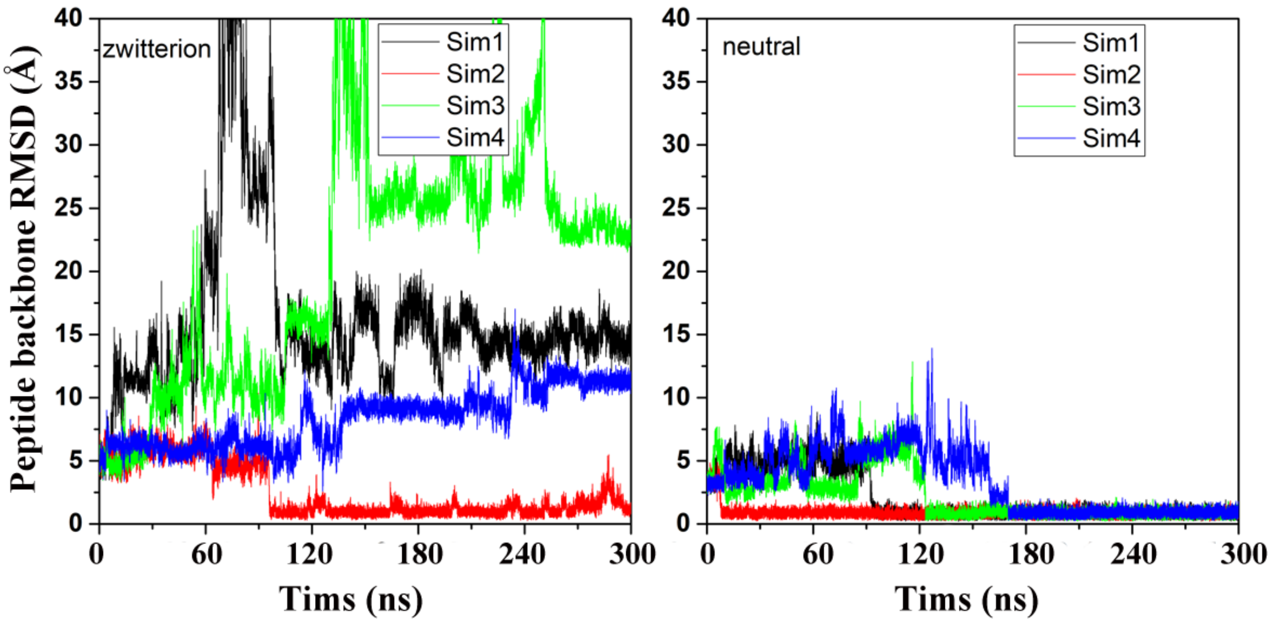


Figure S3. Comparison of zwitterion (left) and neutral (right) terminus models in GaMD simulations of Peptide 2.

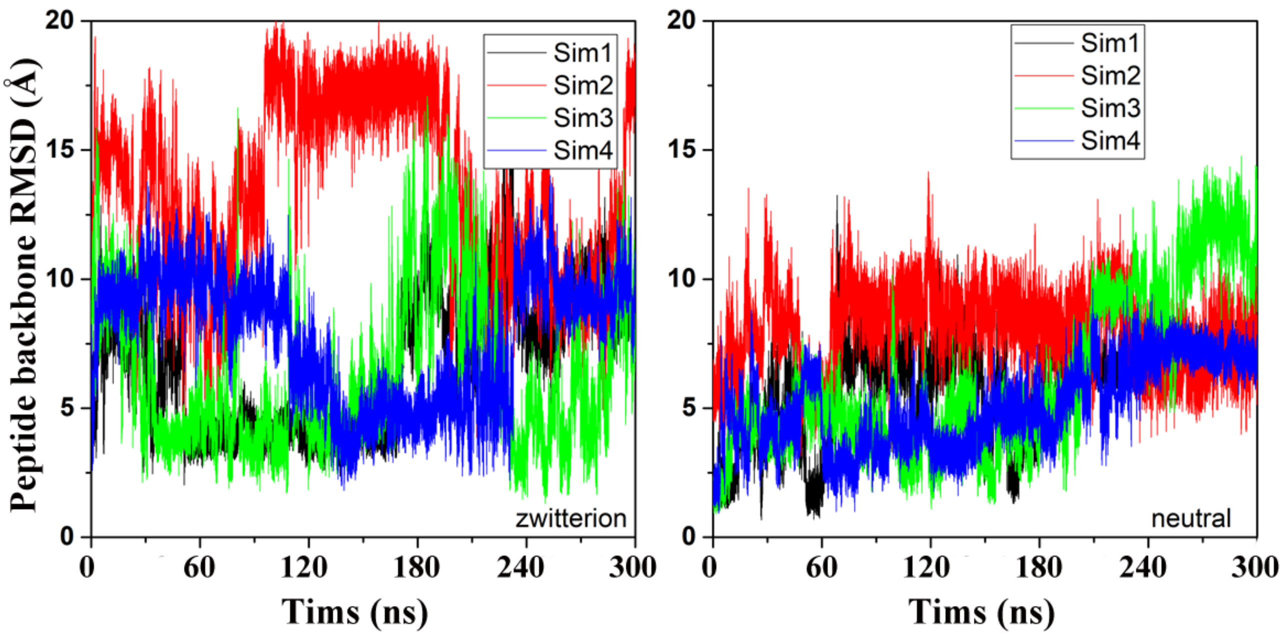


Figure S4. Comparison of zwitterion (left) and neutral (right) terminus models in GaMD simulations of Peptide 3.

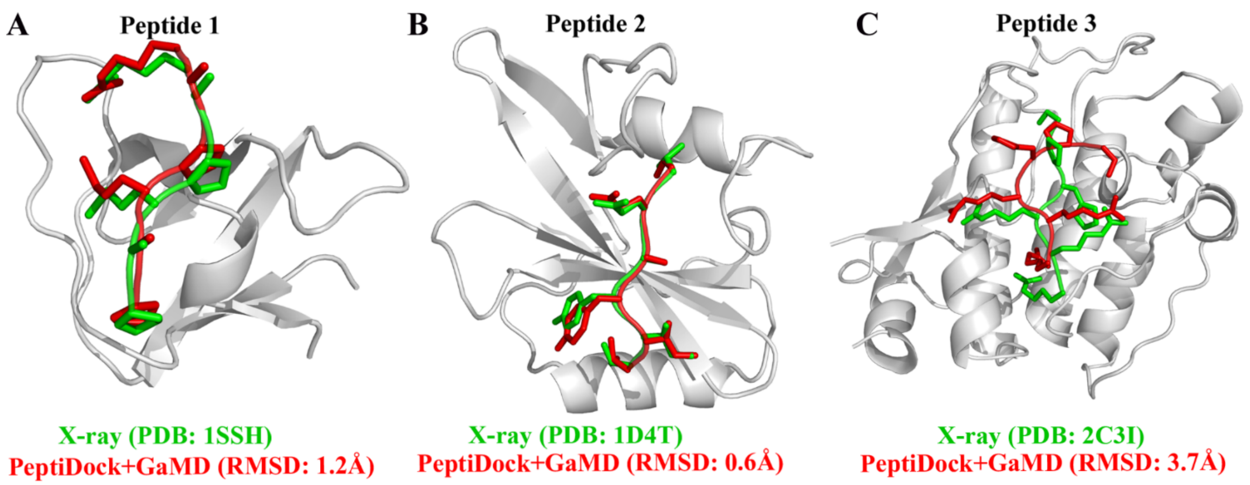


**Figure S5** Binding poses (red) of three model peptides using the zwitterionic terminus models obtained using the "*PeptiDock+GaMD"* are compared with the X-ray structures (green): (A) Peptide 1, (B) Peptide 2 and (C) Peptide 3.
